# Whole genome resequencing of a global collection of Napier grass (*Cenchrus purpureus*) to explore global population structure and QTL governing yield and feed quality traits

**DOI:** 10.1101/2024.10.09.617134

**Authors:** A. Teshome, H. Lire, J. Higgins, T. Olango, E. Habte, A.T. Negawo, M.S. Muktar, Y. Assefa, J.F. Pereira, A.S. Azevedo, J.C. Machado, D.S. Nyamongo, J. Zhang, Y. Qi, W. Anderson, J. De Vega, C.S. Jones

## Abstract

Napier grass (*Cenchrus purpureus*) is a C_4_ perennial grass species native to Sub-Saharan Africa (SSA), primarily used to feed cattle in SSA. In this study, we sequenced the genomes of 450 Napier grass individuals, sourced from 20 different countries. More than 170 million DNA variants (SNPs and Indels) were detected, of which ∼1% informative SNPs were used to assess genetic diversity in the collection. Our resequencing study provided valuable insights into the genetic diversity across a global Napier grass collection. Furthermore, a genome-wide association study on two independent populations, identified multiple quantitative trait loci (QTL) that were significantly associated with desirable agronomic traits, such as biomass yield, nitrogen and cellulose content. Therefore, our results will serve as a valuable resource in safeguarding and unravelling the patterns of Napier grass genetic diversity, in the face of climate change, and spearhead genomics-based breeding programs to develop high-yielding and drought-tolerant varieties suitable for forage and biofuel production.

## Introduction

Globally, grasslands cover 26% of the land area, 70% of agricultural land and play an important role as livestock feed, particularly in Sub-Saharan Africa (SSA)^1^. In SSA, popular grasses include *Cenchrus*, *Urochloa*, and *Megathyrsus* species and these grasses are critically important for smallholders and frequently used by women to maintain the livestock production systems ^2,3^. Unfortunately, annual milk and meat production in SSA remains low compared to the global average^4^. One of the significant reasons behind the below-par productivity of the livestock industry is the inadequate access to quality feeds and forages, worsened recently by the risks associated with climate change^5,6^. Most small-scale livestock farmers in SSA rely heavily on natural common grazing lands as their primary source of forage and feed supply, mainly available during the rainy seasons^7^. Unfortunately, such grazing lands are dwindling because of the inevitable population increase, climate change, and more land being allocated for food crops^8–10^. Consequently, livestock farmers are now more in need of productive, high quality and resilient forage varieties to support their livestock.

Napier grass or Elephant grass [*Cenchrus purpureus* (Schumach.) Morrone syn. *Pennisetum purpureum* Schumach.] is a crucial traditional forage species in SSA, growing mainly up to 2,000 metres above sea level in the tropics^11,12^. It is primarily used to feed cattle in cut and carry feeding systems in Ethiopia, Kenya, Uganda, Tanzania and Nigeria^13–17^ because of its low cost of production, year-round availability under limited irrigation, and some degree of resilience against drought^12,18,19^. Due to its high biomass yield and desirable nutritional traits, Napier grass has recently garnered interest as a candidate for bio-based products and biofuels in tropical and semi-tropical regions of the world, such as the USA and Brazil^20–22^. Once established in the main production field, Napier grass can grow and be maintained for decades under good management practices, yielding up to 50 tons of dry matter ha^-1^ per year^12,23^. Because of its adaptability, persistence, and versatility, it has been naturalized to Central and South America, the tropical parts of Asia, Australia, the Middle East, and the Pacific Islands^14,24^.

Napier grass has the potential to be included in the mainstream feed chain, particularly in the tropics, if research is focused on this species, it can also contribute to energy requirements for current and future generations. Unfortunately, Napier grass remains an underutilized crop with limited genetic and genomic tools developed to date and few cultivars available for farmers. The first reference genome was reported in 2020^25^ and the second and improved one was reported in 2022^26^. The availability of these reference genomes facilitates the generation of molecular markers by elucidating their genomic positions. Here, we report on a species-level whole-genome sequencing (WGS) study of a global collection of 450 genotypes. We analysed the collection’s diversity to explore how breeding, selection and environmental pressures have shaped the Napier grass genome in the international collections. We also analysed the genomic regions associated with important agronomic traits, such as fresh biomass yield and plant height, and nutritional feed-quality traits, such as crude protein content. Thus, the genomic tools developed in this study will enable forage breeders to apply advanced plant breeding procedures such as genomic selection and marker-assisted breeding, which have been lacking to date for Napier grass. Furthermore, new perspectives from the study should benefit conservation efforts worldwide.

## Results

### Phenotypic Variability Among Napier Grass Genotypes

Field evaluations in wet and dry seasons at the Embrapa Dairy Cattle in Brazil indicated significant differences between seasons, among some agronomic and feed-quality traits (Table 1). Plant height (PH) and total fresh weight (TFW) were significantly higher during the wet season whereas cellulose and ash concentrations exhibited no seasonal variation. Furthermore, the interaction between genotypes and harvest cycle was insignificant for most traits, except PH, TFW, dry matter (DM) and total dry weight (TDW). Also, the interaction of genotypes with the season was significant for most traits except TDW, hemi-cellulose content (HCEL) and nitrogen content (NIT), indicating that the performance of genotypes was differentially affected by season. The mean performance per accession for all traits is presented in Supp Table 1.

**Table 1.** Mean square from combined analysis of eleven growth and forage biomass yield traits of Napier genotypes evaluated for two years (2014 and 2016) in Brazil.

| Source of variation | DF | PH | TFW | DM | TDW | CEL | LIG | ADF | NDF | HCEL | IVDDM | NIT | ASH |
| --- | --- | --- | --- | --- | --- | --- | --- | --- | --- | --- | --- | --- | --- |
| Rep | 1 | 4.1*** | 1830** | 0.02*** | 67ns | 79.4*** | 0.01ns | 17.6*** | 4.2ns | 61*** | 148.3*** | 0.55*** | 1.8**<br>* |
| Acc | 90 | 0.83*** | 2036*** | 0.008*** | 289*** | 7.8*** | 3.2*** | 14.2*** | 12.1*** | 4.7*** | 30.7*** | 0.037**<br>* | 3.57*** |
| Harvest | 3 | 41.97*** | 23713*** | 0.2*** | 4360*** | 1636**<br>* | 187.62*** | 2735.3*** | 160.9*** | 1785.6*** | 2575.6** | 0.37*** | 47.65 |
| Season | 1 | 238.19*** | 145149*** | 0.02*** | 4360*** | 1.1ns | 189.4**<br>* | 96.5*** | 323.3*** | 66.5*** | 1581.6**<br>* | 0.3*** | 0.03ns |
| Acc:Harvest | 267 | 0.13*** | 349*** | 0.0018*** | 48*** | 3.2ns | 0.85ns | 5.3ns | 4.3ns | 2.3ns | 7.9ns | 0.007ns | 0.98ns |
| Acc:Season | 89 | 0.23*** | 224* | 0.0035*** | 28ns | 6.2*** | 1.67*** | 11.5*** | 8.1*** | 2.5ns | 14.4*** | 0.01ns | 1.82** |
| Error | 442 | 0.14 | 172 | 0.001 | 23 | 3.6 | 0.76 | 5.8 | 4.1 | 2.1 | 9.1 | 0.01 | 1.08 |

DF – degrees of freedom; PH – plant height; TFW – total fresh weight; DM – dry matter concentration; TDW – total dry weight; CEL – cellulose; LIG – lignin; ADF – acid detergent fibre; NDF – neutral detergent fibre; HCEL – hemicellulose; IVDDM - in vitro dry matter digestibility; NIT – nitrogen; ASH – ash; Acc – accessions

Among the 91 genotypes evaluated, the highest total fresh weight (TFW) was recorded for genotypes BAGCE2, BAGCE64 and BAGCE60. The highest biomass-yielding genotypes as total dry weight (TDW) were similar, indicating a high correlation between TFW and TDW. Regarding nitrogen content (NIT), a key trait in feed quality, the genotypes BAGCE58, BAGCE82 and BAGCE1 were the top performers (Supp. Table 2). Interestingly that genotype BAGCE82, with the highest NIT content, also showed a high mean TFW (64.8 Mg ha^-1^).

The output from the field evaluation trial in Bishoftu, Ethiopia, has previously been reported^12^. Among the shared genotypes, BAGCE53, 86 and 97 performed well regarding TFW in the trials in both Brazil and Ethiopia. Principal component (PCA) and clustering analyses were conducted among the subset of genotypes from Embrapa (Fig. 1). In this analysis divergence was observed based on growth, forage yield and nutritional quality traits. The PCA identified the first three components, explaining 77% of the cumulative variation (Supp. Table 3). The first principal component (PC1) accounted for 40.1% of the total explained variance, PH (0.43), cellulose (CEL) (0.84), acid detergent fibre (ADF) (0.95), and neutral detergent fibre (NDF) (0.78) were the main contributing traits for this component. Likewise, the second principal component (PC2) accounted for 21.1 % of the total explained variance and PH (0.54), TFW (0.89), TDW (0.93), HCEL (0.48) and DM (0.36) were the main contributors for this component (Supp. Table 3).

**Fig. 1.**
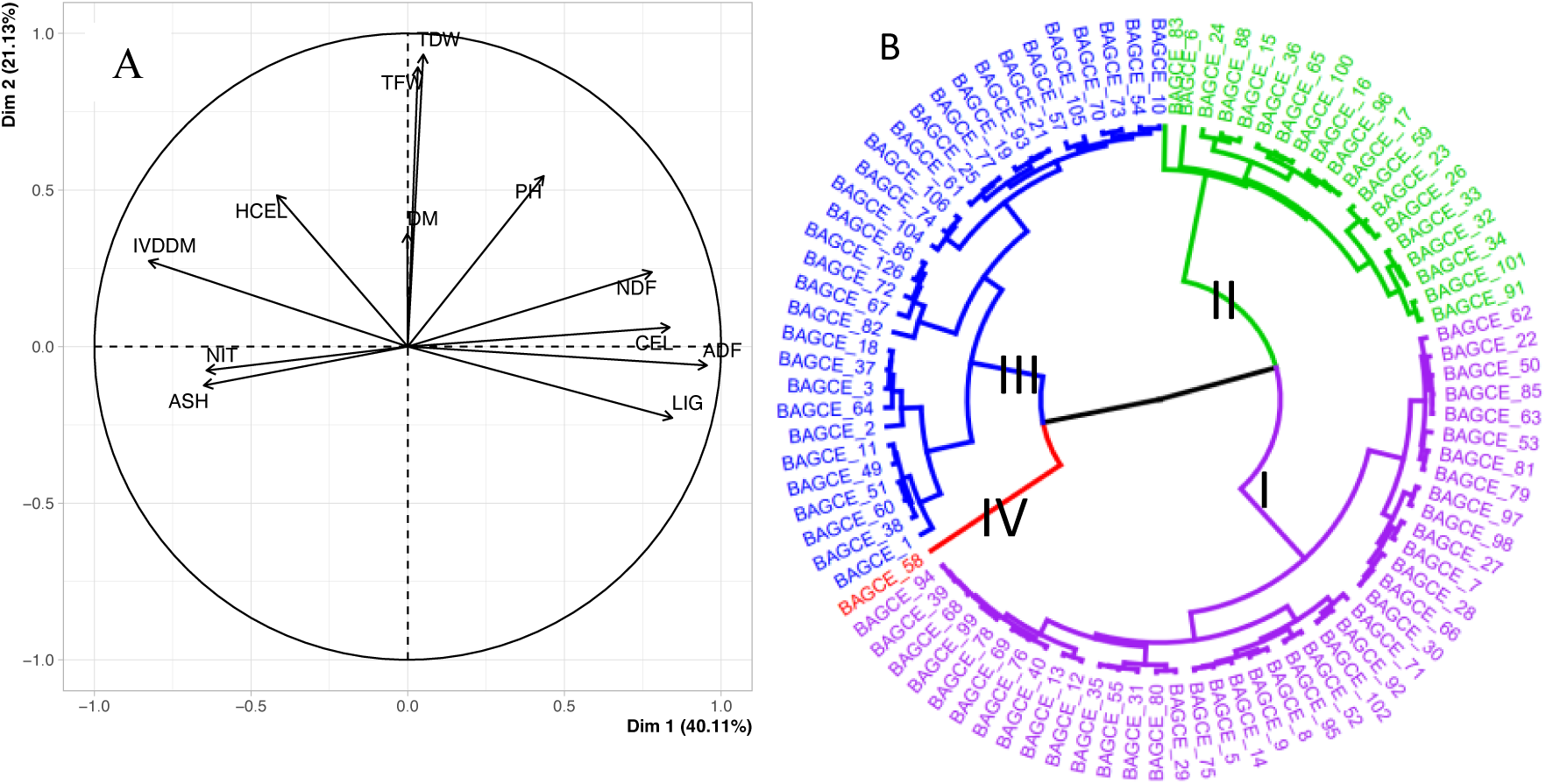
Principal Component (A) and Cluster Analysis (B) based on agro-morphological and nutritional traits among 91 Napier grass genotypes in the Brazil trial

A PCA biplot shows the degree of correlation among measured traits, with those in the same dimension and a tight angle between vectors indicating a high and positive correlation (Fig. 1A). The strongest positive correlation was found between TDW and TFW. Furthermore, CEL and ADF were also highly and positively correlated. TFW, TDW, NDF and PH appeared in the same dimension with a positive correlation with each other, and CEL and ADF were negatively correlated with NIT and ash (Supp. Fig. 1). Based on data from two seasons and five harvest cycles of mean values of 12 quantitative traits, the clustering analysis revealed four major clusters, cluster I, composed of two major sub-clusters, consisted of 40 genotypes; cluster II contained three major sub-clusters and consisted of 19 genotypes; cluster III contained three major sub-clusters and was composed of 31 genotypes; and the cluster IV exhibited a distinct genetic profile, forming an isolated group a single accession (Fig. 1B). Top raking genotypes, in terms of TFW/TDW, such as BAGCE2, BAGCE60, and BAGCE64 were, in cluster III. Interestingly, BAGCE58, which clustered in group IV, by itself, scored the lowest mean TFW and TDW (18.8Mg and 5.9Mg respectively) and the highest NIT content (0.66%).

### Genome-wide SNP Discovery and Their Distribution Across Assembled Chromosomes

Illumina 150-bp paired-end reads were generated from 450 Napier grass genotypes. The average sequencing depth was 15-20x per accession. Nearly ∼170 million variants (SNPs and Indels) were generated and from these variants, ca. 1M hard-filtered SNPs were mapped across the 14 assembled chromosomes of Napier grass. These markers were used for genetic diversity and marker-trait association analyses (<u>Supp.</u> Fig 2). The number of SNPs per chromosome was variable, with more SNPs mapped on the longest A01 and B01 chromosomes (Supp. Table 4). The SNP density was similar for all chromosomes with one SNP detected for every 1,830 bases. We have generated a comprehensive Napier grass genome variation dataset, identifying numerous SNPs from diverse landraces, varieties, and progenies.

### Genetic Variation and Relationship

The PCA revealed an interesting trend with three major clusters and some level of aggregation based on the region of origin (Fig. 2). The first and second coordinates explained 69.9% and 29.9% of the total variation among the genotypes, respectively. Q4 and Q6 groups contained genotypes from the Kiambu and Kakamega districts in Kenya, respectively. However, these two sub-clusters Q4 and Q6 were far apart in the PCA plot. Also, Q9 contained only genotypes from Kenya, although these genotypes originated from three different districts in Kenya (Supp Table 5). Furthermore, Q5 also mainly consisted of genotypes from Kenya and two admixed genotypes sourced from ILRI. There were reportedly eight interspecific hybrid genotypes included in this study (Supp. Table 6) and these hybrids failed to aggregate independently from the rest of the collection. Similarly, genotypes sourced from Embrapa clustered together exclusively in Q8, along with a single accession obtained from USDA. ILRI genotypes were distributed across most clusters, which supports their global origin. The progeny genotypes clustered together mainly in Q10 although some of them were dispersed in different sub-clusters. Interestingly, the admix group which is the largest group with 152 genotypes consisted of germplasm bank genotypes (from Ethiopia, China and Brazil), progenies and breeding lines and this group also contained five of the seven purples genotypes (CN96273, CN96211, CN94131, CN93182 and CN93081).

**Fig. 2.**
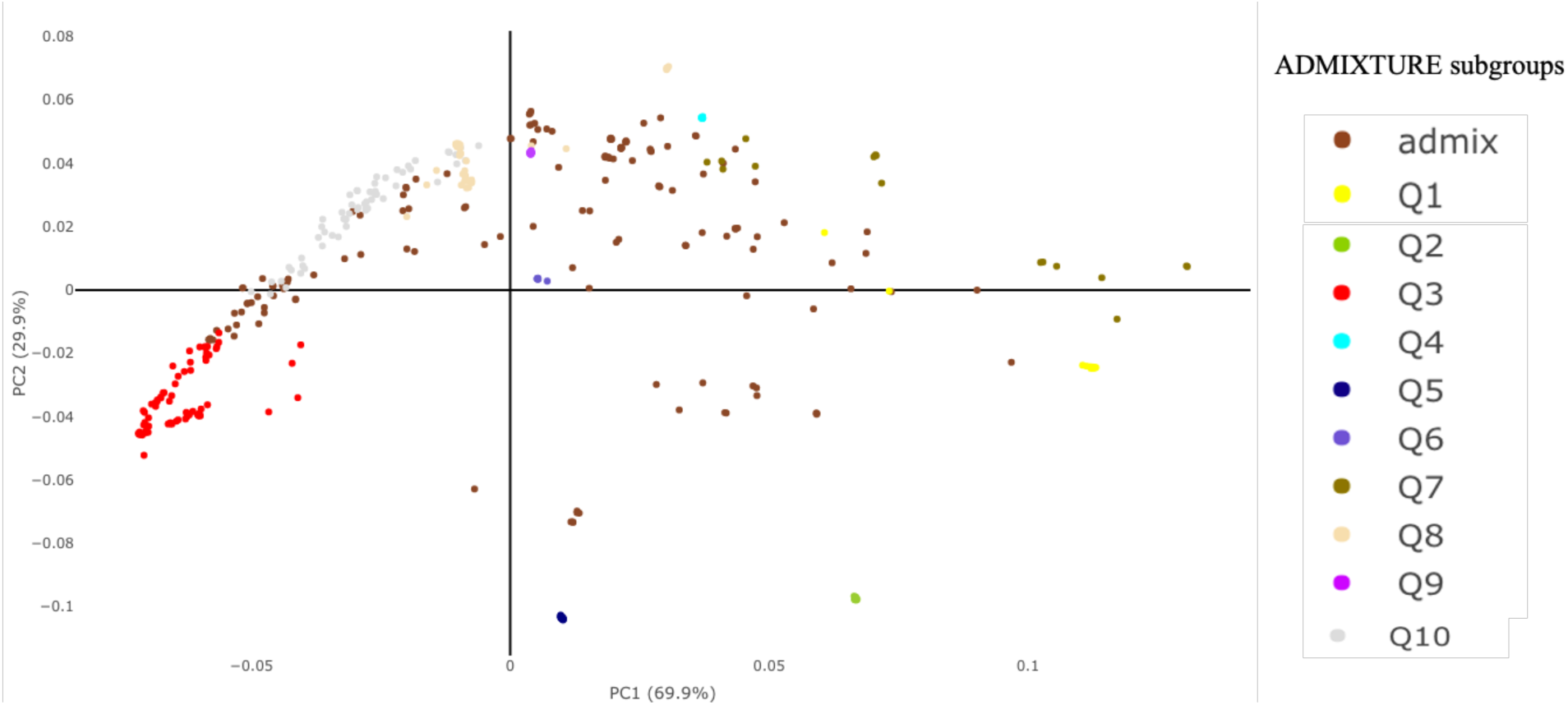
Principal component analysis (PCA) based on ca. 1M SNPs. A scatter plot of PC1 (explaining 69.9% of the variance) versus PC2 (explaining 29.9% of the variance). Colour codes represent the ten sub families defined by ADMIXTURE analysis

### Population Structure Among Global Napier Grass genotypes

Population structure analysis divided the 450 genotypes into ten subgroups according to cross validation (CV) errors (Fig. 3A). Clustering at *K*=6 (Fig. 3B) which separated some genotypes from Embrapa (Brazil), Kenya and ILRI from the rest. However, a high admixture was noticed within the overall collection. A similar trend was observed in the phylogenetic tree, where genotypes were distributed regardless of their region of origin (Fig. 4). For example, two genotypes of Chinese origin (cpReyan4 and PgJujun) clustered together with genotypes sourced from genebanks in Embrapa, Kenya and ILRI. Interestingly, the reference genome CpPurple, a purple variety and all other purple varieties were clustered to each other except BA97. Furthermore, CpPurple showed a high similarity with BAGCE105 (a purple accession from Brazil) indicating that these two could be related genotypes although currently grown on different continents. Progeny genotypes from ILRI showed a mixed trend in the phylogenetic tree; even those originating from the same mother plant did not cluster together. A high level of genetic similarity among Kenyan genotypes was observed indicated possible duplications in the Kenyan collection.

**Fig. 3.**
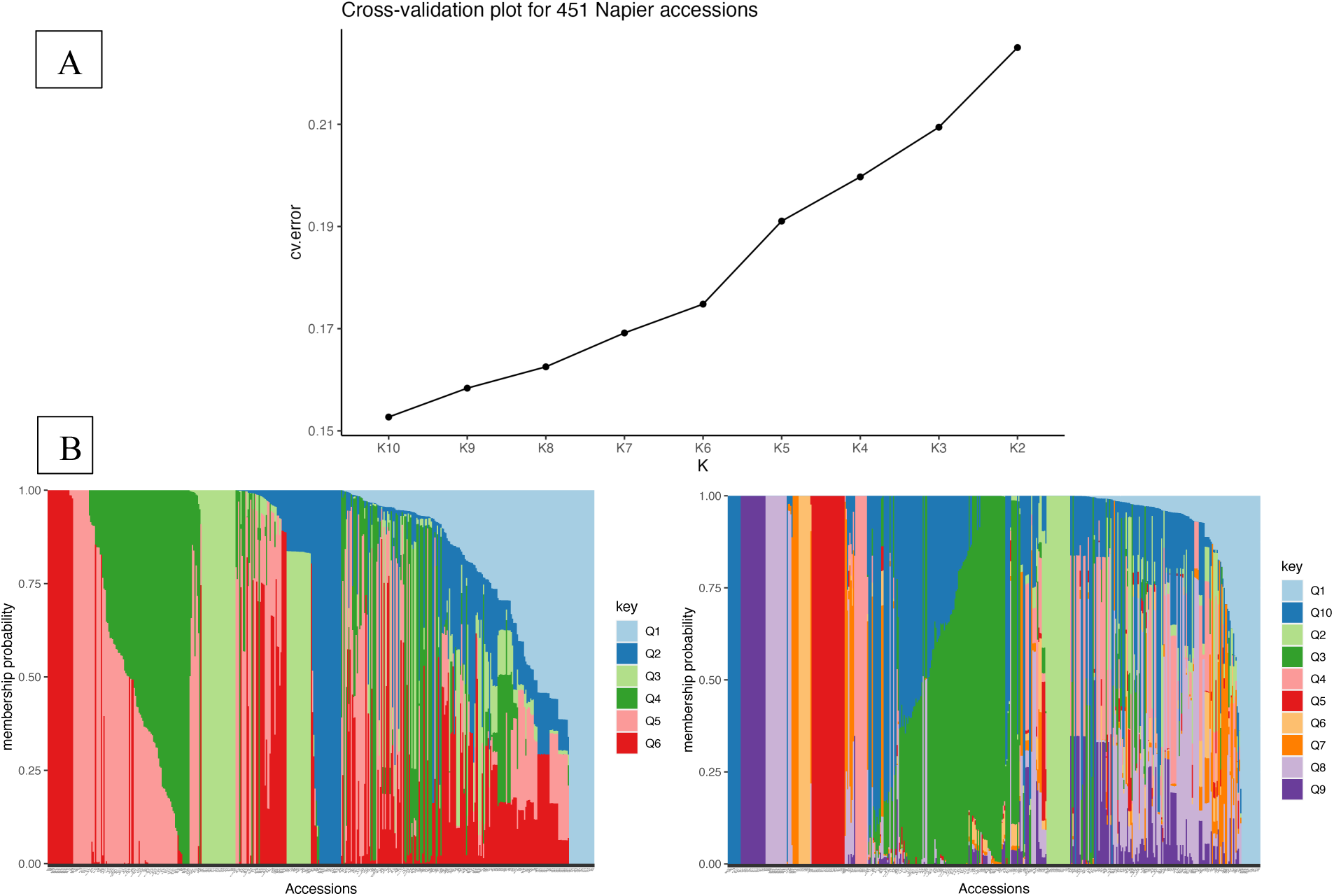
Plot of ADMIXTURE cross validation error from K=2 through K=10 (A). Admixture analysis for 450 Napier grass genotypes results for K=6 and 10 (B)

**Fig. 4.**
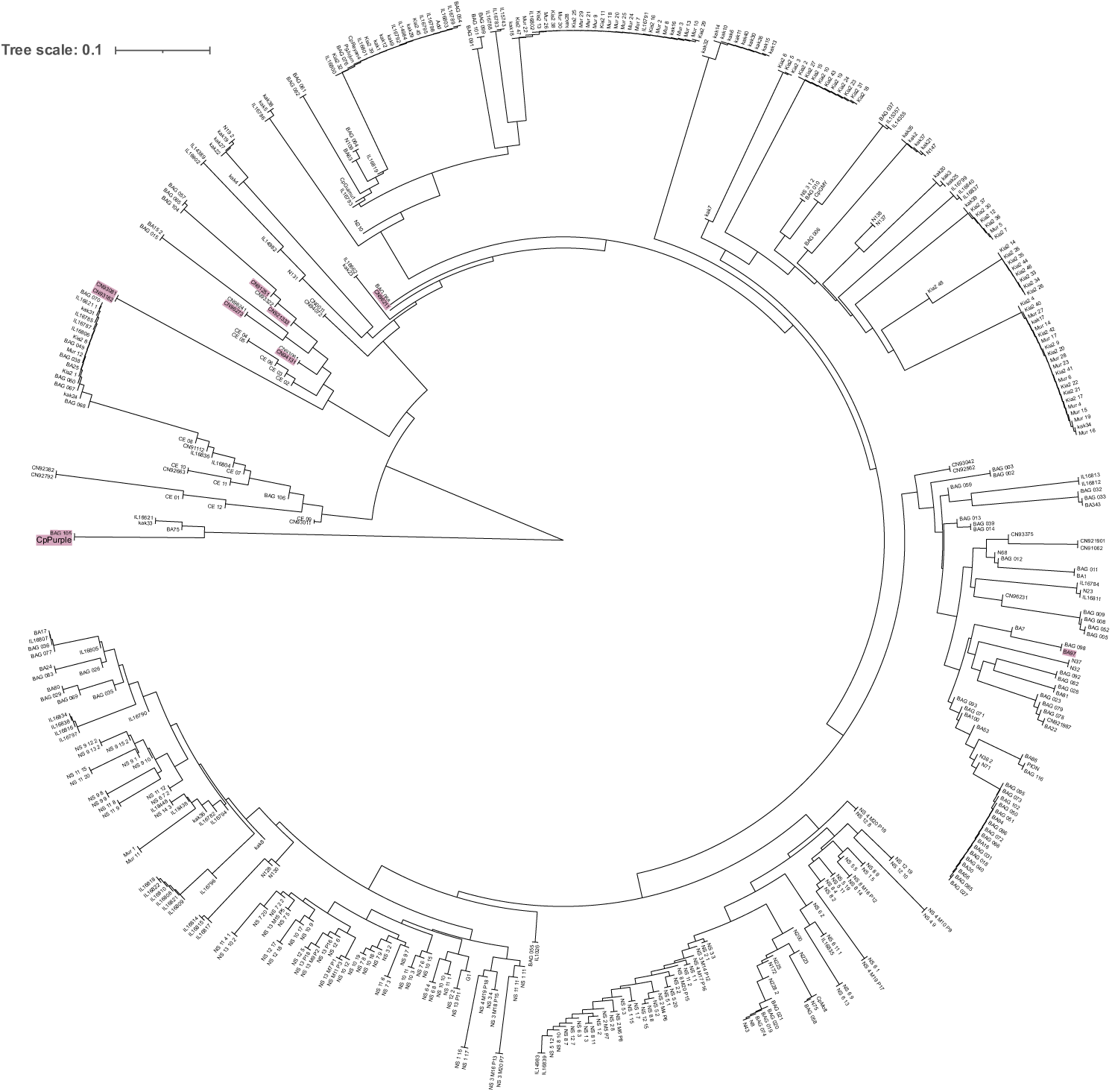
Phylogenetic relationships among 450 Napier grass accessions based on ca. 1M filtered SNPs. Accessions with purple background are the ones with purple leaf colour and the reference accession, CpPurple, was one among the purple accessions

### Marker-trait associations

For the field evaluation carried out in Brazil, the marker-trait association analysis was carried out independently for dry and wet seasons for each of the 12 quantitative traits (Supp. Fig. 3). Significantly associated SNP locus (−log10(p) < 5), were identified for ten traits scored in more than one association model for either of the seasons (Supp. Fig. 4). Interestingly, significant QTL were recorded for some traits such as TFW and ADF (> −log10(p)) value of 5) mainly in dry season (Supp. Table 7).

A genome-wide association study (GWAS) was also carried out for the trial conducted in Ethiopia, and the results revealed interesting associations for the nine traits scored. All traits significantly associated with QTL were identified under dry and wet seasons and also under two soil moisture conditions. QTL significantly associated with stem thickness (ST) was identified during the wet season and under both soil conditions in the dry season (Supp. Table 8). Likewise, SNPs were identified to be significantly associated with total fresh weight (TFW) under both dry and wet seasons, in both soil conditions. For the binary trait, leaf colour (green vs. purple), a total of 494 SNP loci were determined to be significantly associated (8 < −log10(p)) using three different GAPIT models (Supp. Table 9). In general, a total of 207 SNP loci were significantly associated with other traits, excluding the leaf colour, using three GAPIT models, for the trial carried out in Ethiopia (Supp. Fig. 5). Forty-seven of those marker trait associations (MTAs) were significantly associated with more than one trait and were also significant in more than one GAPIT model. Three traits, PH, TFW and TDW were measured in both field trials in Ethiopia and Brazil, and the combined data were used to carry out a GWAS analysis that led to the identification of additional QTL (Supp. Fig. 6). Interestingly, significant QTL were identified for all three traits at a higher threshold (7 > −log10(p) in the dry season. In contrast, a significant QTL was only identified for total dry weight (TDW) in the wet season.

From the significantly associated QTLs, thirteen were associated with multiple phenotypic and feed quality traits with more than one GAPIT model. The search for those significant MTAs has revealed sequence similarity with proteins of various functions in the Phytozome database. For instance, a region around a QTL associated with TDW and TFW showed sequence similarity with a Gag-Pol-related retrotransposon. Similarly, another QTL significantly associated with traits such as ADF, *in vitro* dry matter digestibility (IVDMD), and LIG showed similarity with a WDSAM1 protein (Supp. Table 10). Additionally, a QTL highly correlated with leaf colour exhibited similarity with the Zinc-finger domain of a monoamine-oxidase A repressor R1 and carotenoid synthesis regulator regions.

## Discussion

More research is needed on tropical forages compared to temperate counterparts like perennial ryegrass and alfalfa. As a result, only landraces are available for use by small-scale farmers, in most parts of the globe, particularly in Africa, and these may need to be better adapted to current and future climate conditions^3^. Among commonly grown/cultivated tropical grasses, Napier or Elephant grass is the most popular in SSA because of its high biomass production, resilience against high temperatures and water scarcity, and regrowth capability that allows for four to six cuts per year^27,28^.

Two independent collections of Napier grass genotypes, from ILRI and Embrapa, were evaluated in field in Ethiopia and Brazil, respectively. These evaluations were critical to assess phenotypic performance and diversity and to identify the “best bet” candidates for a new cycle of breeding. Breeding aims to increase biomass, resilience against drought stress and high feed quality traits such as high crude protein and low lignin content. Higher values were scored for traits such as PH, TFW and TDW, during the wet rather than the dry season in both Ethiopia and Brazil and a similar finding was also reported previously1^2,29^. On the contrary, feed quality traits, such as cellulose (CEL) and ash content, remained the same regardless of the season (Table 1). Overall, genotypes such as BAGCE2, BAGCE64 and BAGCE60 scored the highest TFW-TDW, and the same genotypes also performed consistently well in both dry and wet seasons, implying their genetic potential. BAGCE30, one of the nine genotypes shared between the trials in Brazil and Ethiopia performed relatively well in the trial in Brazil, with 68.6 Mg ha^-1^ and 40.1 Mg ha^-1^, mean TFW and TDW, respectively. This genotype also showed resilience against drought stress. It produced consistently high total dry biomass under two low soil moisture conditions in the trial in Ethiopia, indicating its genetic potential to perform across geographies^28^.

Feed quality traits such as crude protein, standardly calculated as % nitrogen x 6.25, and lignin contents are key forage traits. In the trial carried out in Brazil, the highest NIT was recorded for accession BAGCE58, but this accession was ranked the lowest for TFW and TDW (Supp. Table 2). Positive correlations were observed between some of the measured traits; for example, TFW and TDW, CEL and ADF were highly correlated (Supp. Fig. 1) and this result is consistent with reports made by ref.28. On the other hand, the traits CEL and ADF were found to be negatively correlated with NIT and ASH.

Napier grass is one of the lignocellulosic crops that have become attractive for bioenergy production due to its high cellulose, hemicellulose and lignin contents and perennial nature^30,31^. This study’s average cellulose (CEL) contents for all genotypes ranged from 36.9 to 43.1%. Hence, genotypes BAGCE83, BAGCE6 and BAGCE59, with higher cellulose content can be targeted for future development of Napier grass varieties for ethanol production. On the contrary, Pioneiro, a released cultivar in Brazil as a livestock forage, was among the genotypes with the lowest cellulose content. A few genotypes, BAGCE104, BAGCE106 and BAGCE62, scored lower for cellulose content than Pioneiro, which implies opportunities for improving digestibility by breeding strategies. Clustering of genotypes based on agro-morphological and feed quality traits grouped the genotypes into four major clusters (Fig. 1). This phenotypic diversity information is valuable, along with genomic tools, in future selection of genetically diverse, phenotypically superior parental genotypes for hybrid breeding. Cluster IV, represented only by BAGCE58, scored the lowest overall mean TFW and TDW (18.8 and 5.9 Mg ha^-1^, respectively) and the highest NIT content (0.66%). On the contrary, high-yielding genotypes such as BAGCE2, BAGCE60 and BAGCE64, are clustered in cluster III. BAGCE58 could improve nitrogen production associated with other parentals and produce output in a breeding program.

Previous genotyping by sequencing (GBS) studies on Napier grass reported ca. 100K SNP markers^19,32–34^ and SNP density/chromosome was low compared with the present study. Whole-genome sequencing of 450 global Napier grass global genotypes, predominantly landraces, generated an unprecedented resource, more than 100 million variants (both indels and SNPs), highlighting the available diversity within this global collection. In the present study, a SNP was detected on average every 1803 bases on all the chromosomes (Supp. Table 4). These genome-wide SNPs can be used as a DNA fingerprinting tool in the germplasm bank collections and to verify the trueness-to-type of cultivars. After a complex filtering, nearly a million SNPs were retained, evenly distributed along the 14 Napier grass chromosomes. Napier grass is an allotetraploid (2n = 4x = 28, A’A’BB sub-genomes), the A’ sub-genome shows a high degree of homology with the pearl millet A genome (*Cenchrus americanus*, 2n = 2x = 14, AA) and the two species can readily hybridise to generate sterile triploid hybrids (2n = 3x = 21, AA’B genome)^35^. Therefore, the genomic tools developed in this study could potentially be used for pearl millet genetic improvement or during the development of interspecific hybrids from these species. Ref.32 generated more than five thousand simple sequence repeat (SSR) markers for Napier grass, following a GBS approach, and from the high SNP rate per chromosome developed in this study, a significantly higher number of SSRs can be created.

Using the filtered SNPs, principal component analysis (PCA) showed three major aggregations (Fig. 2), but this clustering was not aligned with region of origin. This could be because Napier grass originated from Africa, and although the materials used in this study were sourced from 20 different countries, limited natural or artificial genetic selection has taken place in those samples. A similar finding was reported by ref.34, following GBS genotyping and by ref.36 using, AFLP markers, where Napier grass genotypes from different countries did not cluster based on region of origin. However, some genotypes have shown aggregation based on area of origin; for example, samples labelled Q4 and Q6 represent only Kenyan samples, but these samples were from different districts (Kiambu and Murangu). This is not unexpected due to the existence of historical and current exchange of root splits through the informal seed system involving Kenya’s regional and countrywide farming communities. A similar finding was previously reported by ref.34.

Another interesting finding from the PCA was that those progenies arising from 14 mother plants aggregated separately from each other and did not show any unique profile that reflected their sexual origin. Ref.34 reported a similar finding, using GBS genotyping, where progenies showed irregular clustering and did not aggregate with their respective mother plants. Several ILRI genotypes aggregated closer to the Embrapa elite breeding lines that led to the development of cultivars such as BRS Capiacu and BRS Kurumi^37^ which implies their genetic potential for further improvement of released cultivars. In this study, eight hybrids (*Cenchrus purpureus* × *Cenchrus americanus*) were present. Still, these hybrids failed to cluster independently, implying the an error during the acquisition or management of these hybrids. However, before any strong conclusions can be drawn, taxonomic and/or cytology characterization should be performed to confirm their hybrid status.

Population structure analysis was conducted to develop a better understanding of the relationship among genotypes, landraces, breeding lines, progeny plant etc. Population structure analysis divided the 450 genotypes into ten subgroups according to cross validation error values from an ADMIXTURE analysis (Fig. 3B). This analysis separated some of the Embrapa genotypes from the ILRI genebank materials (Supp. Table 5). Ref.34 reported a similar trend, delineating Embrapa and ILRI collections using a GBS approach. Another study by ref.38, using SSR markers, also observed two sub-populations in nearly two hundred Napier grass genotypes from the same collections. Therefore, this result implies that the ILRI and Embrapa collections can be separated into two independent gene pools with slight admixture in between, suggesting that heterotic breeding for desirable traits would be practicle. Interestingly, most progenies were distributed among different sub-clusters which was also the case during the PCA analysis. This could be due to the recombination of genes during hybridization and a similar unorthodox clustering of progenies, which was also reported by ref.34.

A Phylogenetic tree of the 450 genotypes revealed a similar result to that reported earlier, where there was no clustering based on region of origin. In this phylogenetic tree, two major clusters were observed; one cluster consisted of five genotypes and all of the remaining samples grouped in the second cluster. The first cluster included the genotype used for the reference genome (CpPurple) and the rest were from different sources (two from Embrapa, one accession each from Kenya and from ILRI). Interestingly, CpPurple (the reference genome and purple variety) showed high genetic similarity with BAGCE105 (purple genotype), implying these two genotypes could be highly related though sourced from two different countries, China and Brazil respectively. Likewise, Pioneiro and BAGCE116 showed high genetic similarity in the tree, and the accession BAGCE116 was selected from a different elephant grass population than the one in which Pioneiro was present. BAGCE116 is phenotypically like Pioneiro (plant biotype). Still, it exhibits yellowish and greenish stripes on the leaves, providing an ornamental characteristic. Although this has not been conclusively proven, we suggest that it is highly likely to be a mutant plant selected from within the Pioneiro cultivar.

As was observed in the PCA, genotypes labelled as hybrids (IL16835, IL16837, IL16834 IL16838 IL15357, IL16840 and IL14982) did not cluster together, in the phylogeny tree, to reflect their hybrid profile. The present finding provides corroborating evidence to a similar observation made by ref.33. A high level of genetic similarity, and/or possible duplications was observed among genotypes sourced from Kenya. Ref.34 also reported that there is low genetic diversity among Kenyan genotypes despite these genotypes representing collections from different districts in Kenya.

Self-incompatibility^39,40^ and an obligate outcrossing nature ref.25 have been reported in Napier grass, and the phylogenetic tree generated in this study can play a role in selecting distantly related parents to achieve hybrid vigour in Napier grass crosses. Hybrid breeding has been successful in many temperate forages such as ryegrass^41,42^, and a similar approach could benefit Napier grass improvement as it can be propagated by vegetative means and target traits could easily be fixed at the F_1_ stage. In general, the phylogeny has reaffirmed the trend shown by the PCA and ADMIXTURE analyses, that there is a rich diversity in this collection of Napier grass genotypes with no solid evidence to support region of origin clustering.

Though resilient against multiple biotic and abiotic stresses, Napier grass production in SSA is currently being challenged by two main diseases, head smut and stunt, recurrent droughts and feed quality issues among farmers. Developing high-yielding, nutritious and resilient varieties is important to have maximum impact on animal performance. However, field characterization of Napier grass is time consuming and laborious, mainly due to its perennial nature^12^, architecture (big unlike most grass species and hence not suited for greenhouse experiments^43^) and reproduction as obligate outcrossing nature^39^. Hence, developing molecular markers is critical to fast-track the development and release of improved cultivars while saving time and resources. Simulation studies on wheat highlighted that the breeding programs that use genomic assisted selection strategies improved yield at a significantly more rate than those that relied solely on phenotypic selection^44^. Few genome wide associated studies (GWAS) have been performed on Napier grass^19,22,33^. Hence, the high-density and genome-wide SNP markers reported in the present study will significantly enhance our ability to discover markers in linkage disequilibrium with complex traits, such as yield traits (total fresh and dry weight (TFW and TDW) and water use efficiency (WUE)). In the present study, a GWAS was carried out on the two independent collections and interesting marker trait associations (MTAs) were identified. For the field evaluation carried out in Brazil, the GWAS analysis identified SNPs (5 < −log10(p)) significantly associated with all measured traits, using at least one GAPIT model, during both seasons. All in all, for the trial conducted in Brazil, a total of 318 SNPs were significantly associated with 12 traits under both dry and wet seasons conditions (Supp. Table 7) and these MTAs could play a role in future Napier grass breeding programs in Brazil and beyond. The associated SNPs can be also used to search for the genes underpinning important traits and led to the identification of targets for gene editing, a vital tool to increase animal production in the tropics^45^.

For the trial carried out in Ethiopia, a relatively higher number of QTL were detected for traits such as TFW, TDW, and tiller number (TN) which are key agronomic traits for Napier grass improvement. All traits showed significant associations under dry and wet conditions and also under two soil moisture stress conditions (Supp. Table 8). Ref.19 identified QTL that were significantly associated with yield, water use efficiency and feed quality traits in Napier grass. For example, for TDW, dry season and severe water stress (SWS) conditions, significant MTAs were reported on Chr5, Chr9 and Chr13^19^ and in this study more MTAs were identified in most of the Napier grass chromosomes (Supp. Table 8). Traits such as stem thickness (ST) recorded significant QTL (> −log10(p) value of 5) markers during the wet season harvests, and under both MWS and SWS soil moisture conditions during the dry season (Supp. Fig. 5). One of the critical traits scored in this study was leaf colour (green vs. purple) and the GWAS analysis identified 494 highly associated SNPs, distributed along all chromosomes, (8 > −log10(p)) for this binary trait (Supp. Table 9). Ref.19 reported GBS markers highly associated with purple leaf colour in Napier grass, and in the present study, a significantly higher number of SNPs were identified in both sub-genomes. Purple colouration in Napier grass is due to high anthocyanins content which has potential health benefits to both humans and animals^46,47^. Hence, the high number of MTAs identified for this trait will be useful in future Napier grass feed quality improvement. Three traits, PH, TFW, and TDW, were measured in both field trials in Ethiopia and Brazil, and the combined data were used to identify QTL that were consistently found across environments. Interestingly, in the dry season treatments, significant QTL were identified for all three traits at a higher threshold (7 < −log10(p). Still, in the wet season, a significant QTL was identified for TDW only (Supp. Fig 5).

The region around one of the significantly associated SNPs for leaf colour (A01_63825491) showed synteny with homologs responsible for protein-serine/threonine kinases (Supp. Table 10). This family of proteins act as a central processor unit receiving input signals from receptors capable of sensing environmental and other external cues translating them into appropriate information that triggers changes in gene expression and metabolism which ultimately manipulate various functions such as growth and development, fertilization, and immunity^48–50^. Another SNP (A01_ 63717178) significantly associated with leaf colour was located in the region that showed synteny with genes encoding proteins for carotenoid pathways in grass species such as rice. The carotenoid biosynthesis pathway plays a key role in producing photosynthetic and protective pigments, stress hormones and various volatiles^51,52^, and these secondary metabolites involve in multifaceted physiological functions in a myriad of species including humans and animals. One of these functions is antioxidant properties, mitigating diseases incidence in both plants and animals^53^. Hence the markers identified in the present study could be utilized to develop nutritionally superior Napier grass cultivars in the future.

## Conclusions

Limited access to high quality forages is one of the major factors affecting livestock performance in SSA, and the use of indigenous forage species such as Napier grass is recommended as these forages are familiar to smallholder farmers, require low inputs and are adaptable to different agro-ecologies and production systems. Our results highlight genomic differences and marker trait associations in global Napier grass genotypes, varieties and landraces, which are likely the product of adaptation to multiple environmental conditions and breeding. We hope that this genomic tool (diversity profile for the global collection and QTL identified) and recently available reference genomes will encourage the routine use of molecular markers for Napier grass improvement programs. Furthermore, these molecular and genomic resources are crucial for managing genetic resources and advancing conservation programs for Napier grass, both in farmers’ fields (*in-situ*) and in genebanks (*ex-situ*).

## Materials and Methods

### Napier Grass Field Evaluation

Phenotype data were assessed from two collections of Napier grass genotypes which were independently evaluated in this study. The first collection consisted of 84 genotypes conserved at the International Livestock Research Institute (ILRI) genebank which was evaluated in Bishoftu, Ethiopia for two consecutive years, in P-rep design, replicated twice. Details of the field evaluation have previously been reported^12,19^. Briefly, the genotypes were exposed to different soil moisture stress conditions namely rainfed (RF) wet season conditions (approximately 30% volumetric soil water content -VWC), moderate water stress (MWS - with 20% VWC), and severe water stress (SWS - with 10 % (VWC)) during the dry seasons, over two years and multiple cuttings. The second collection of 91 Napier grass genotypes was evaluated at the Embrapa Dairy Cattle Research Centre experimental field, located in Brazil, and five cuttings were conducted between 2014-2016 in both wet and dry seasons. Ref.22 described in detail these evaluations that were done in natural conditions. Nine genotypes (BA17, BA30, BA34, BA53, BA81, BA86, BA93, BA97 and Pioneiro (released cultivar) were shared between these two trials.

### Phenotyping of Agronomic and Feed Quality Traits

The following agronomic traits were measured for the trial carried out in Ethiopia; leaf length (LL, mm), leaf width (LW, mm), leaf to stem ratio (LSR), stem thickness (ST, mm), tiller number (TN) and biomass yield data (total fresh weight (TFW, g) and total dry weight (TDW, g) were collected as described previously in^28^. Water use efficiency (WUE) was also calculated by dividing the total dry weight per plant by the total volume of irrigated water applied to each plant during the dry season. Likewise in Brazil, plant height (PH, m), production of total fresh weight (TFW, Mg ha^-1^), production of total dry weight (TDW, Mg ha^-1^) and dry matter concentration (DM, %) were scored. Furthermore, nine feed-quality nutritional traits including acid detergent fibre (ADF), neutral detergent fibre (NDF), lignin (LIG), cellulose (CEL), hemicellulose (HCEL), *in vitro* dry matter digestibility (IVDMD), ash (ASH), and nitrogen content (NIT) were also scored in Brazil. The DM concentration recorded for agronomic traits was used as a common denominator for estimating of biomass digestibility. Further details can be found in^22^.

### Phenotypic Data Analysis

Collected phenotypic values for each trait were checked for normal distribution and transformed, when needed, ahead of variance comparison using the bestNormalize R package^54^. Phenotypic variability between genotypes was calculated with R statistical software^55^ using the model: E · R + E · R · B + G + G · E where E, R, B, and G denote environment, replicate, incomplete block, and genotype, respectively. Environment effects and replicate effects nested within the environment, both represented by (E.R), were taken as fixed. In contrast, the block effect nested within the replicate and environment and the genotype-by-environment interaction (G·E) were taken as random. The main genotype effect (G) was taken as random. Analysis of variance (ANOVA) and multiple comparison tests (LSD) were conducted at a probability level of 5%. Furthermore, phenotypic data were used to carry out hierarchical clustering and principal component analysis (PCA) using the Factoextra R package^56^. Correlation analysis was also carried out between all the variables measured in the field evaluation in Brazil.

### Sequenced Worldwide Napier Grass Collection

A total of 450 Napier grass genotypes were sequenced and deposited as bioproject PRJEB73794: 62 from the ILRI genebank, 131 from Embrapa, 23 from the USDA, six from China (Lanzhou University), 118 from the Kenya Agricultural and Livestock Research Organization (KALRO) and two released cultivars, namely Super Napier (G1) and Pioneiro (PION)). In addition, 109 progeny plants (generated from seeds collected from 14 ILRI genotypes (mother plants) by open pollination were sequenced. The progenies were from open pollinated in the field and the pollen donor genotypes were unknown. All mother plants were represented by 6-10 progenies represented all mother plants except for one mother plant (IL18438), which a single progeny represented. More information about these genotypes can be found in Supp Table 1: metadata.

### DNA Extraction and Sequencing

Young leaf tissue was collected from respective genotypes and subjected to isolation of genomic DNA following the procedure described in the Qiagen DNeasy® Plant Mini kit (250) (Qiagen Inc., Valencia, CA). Before library preparation, DNA quality was checked on 1% agarose gels, and DNA purity was checked using a Nanophotometer® spectrophotometer (IMPLEN, CA, The USA), and DNA concentration was measured using the Qubit® DNA Assay Kit in a Qubit® 2.0 Fluorometer (Life Technologies, CA, USA). High-quality DNA with a minimum of 50 ng/µl was used for Illumina whole-genome sequencing. The genotypes were sequenced using Illumina technology using paired-end 2 × 150 bp short-reads. A total of 4.92 Tb of data was generated, with an average sequencing depth of 15-20x per sample. Library preparation and sequencing were conducted by Novogene (https://en.novogene.com).

### Read Mapping, SNP Calling and Filtering

The quality of raw reads was checked using the FastQC^57^ and MultiQc tools^58^. Afterwards, raw reads were trimmed and filtered with the trimmomatic tool^59^ to remove Illumina Truseq adapter remnant sequences, as well as low-quality reads (with a quality score lower than 30). Curated reads were mapped against the Napier grass reference genome^25^ with the Burrows Wheller Aligner (BWA)^60^. The SAM files generated from the BWA step were converted into sorted BAM files using SAMtools^61^. The HaplotypeCaller tool, Genome Analysis Toolkit (GATK4.4), was used for the variant calling step with default parameters^62^. The generated vcf file, from the variant calling step, was filtered and pruned with BCFtools (v.1.9)^61^. The SNP filtering step retained SNP loci that were biallelic, polymorphic, read depth between 10 and 300, mapping quality (GQ>20), with minor allele frequency above 0.05 and missing calls in less than 1% of the samples.

### Genetic Diversity and Population Structure

The population structure analysis tool, ADMIXTURE (v1.3.0)^63^, was used to infer optimal cluster/subpopulations (*K*) and the proportion of ancestry among the 450 global Napier grass genotypes, with a thinned set of 1,068,685 SNPs. Ten independent runs were carried out for maximum likelihood estimates of the ancestry subgroups (*K*) from 2 to 10. For each *K*, ADMIXTURE was run 20 times with varying random seeds. Afterwards, CLUMPP software^64^ was used to align up to 10 Q-matrices in the same cluster. The number of ancestors was determined according to the position of the minimum value, with an error rate obtained from the cross-validation (CV) score. A good value of *K* will exhibit a low cross-validation error compared to other *K* values. Outputs from ADMIXTURE were collated using the R pophelper program (v.2.3.1)^65^, which compares the ancestral make-up of each predicted population.

Using genotypic data, principal component analysis (PCA) was performed to examine inter-population distribution using the SNPRelate (v. 4.0.2)^66^ and Plotly R packages^67^. A phylogenetic tree was also constructed with the filtered, high-quality SNPs using identity by descent (IBD) with the SNPRelate R package (v.4.0.2)^66^ and visualized with the interactive Tree Of Life (iTOL)^68^.

### Genome-Wide Association Study (GWAS)

A total of 174 genotypes in two independent populations, 90 from Embrapa and 84 from ILRI, were considered for the Genome-Wide Association Study (GWAS). These genotypes included two independent populations phenotyped in the field evaluation carried out in Brazil and Ethiopia. These trials were carried out at two different times, and different traits were measured in each experiment. For the field evaluation carried out in Brazil, the marker-trait association analysis was carried out separately for dry and wet seasons, for each of the 12 quantitative agronomic and feed-quality traits (plant height - PH, total fresh weight - TFW, dry matter concentration - DM, total dry weight -TDW, cellulose - CEL, lignin - LIG, acid detergent fibre - ADF, neutral detergent fibre - NDF, hemicellulose - HCEL, *in vitro* dry matter digestibility - IVDDM, nitrogen - NIT and ash - ASH). In the experiment carried out in Ethiopia, the traits measured were plant height (PH), leaf length (LL), leaf width (LW), leaf to stem ratio (LSR), stem thickness (ST), total dry weight (TDW), total fresh weight (TFW), tiller number (TN) and water use efficiency (WUE) and the groups were split into moderate water stress (MWS) and severe water stress (SWS) dry season treatments.

Furthermore, GWAS was employed to investigate marker-trait associations for three agronomic traits; plant height (PH), production of green biomass (TFW), and production of dry biomass (TDW) assessed in both field evaluations conducted in Brazil and Ethiopia. The mean value for each trait was first indexed as low, medium and high based on quantile values for each trait per season, i.e., mean values less than the 1^st^ quantile were labelled as low and mean values between the 1^st^ and 3^rd^ quantiles were labelled as medium and mean values above the 3^rd^ quantile were labelled as high. Once recoded, the data from each country was merged per trait and for each season. The transformed data from the nine genotypes, shared between the two evaluations showed some degree of inconsistency (probably due to GxE interactions), particularly for the dry season and in such cases data from Ethiopia was selected since frequent measurements, i.e. every eight weeks, were taken during the trial in Ethiopia (Supp Table 1).

The average values for all traits were normalized ahead of the GWAS using the bestNormalize R package^54^. Different GWAS models were used, Fixed and random model Circulating Probability Unification (FarmCPU)^69^, Bayesian-information and Linkage-disequilibrium Iteratively Nested Keyway (BLINK) model^70^ and the Multiple-Locus Mixed Linear Model (MLMM)^71^ implemented in the Genomic Association and Prediction Integrated Tool (GAPIT) version 3 software package within the R environment^72^. Population structure was accounted for by including 3-5 principal components in each GWAS models applied to improve marker-trait association prediction. The distribution of observed vs. expected −log10(p) values was visualized using Quantile–Quantile (Q–Q) plots, graphical representations of the deviation of the observed *p-*values from the null hypothesis. In this study, a minimum threshold *p*-value of 0.00001 (–log10(p) > 5) was used to declare significant SNPs linked with target traits.

Regions of 0.04Mbp around highly significant SNPs (identified by multiple models and associated with more than one trait and/or treatment condition) were blasted against protein databases such as Phytozome databases^72^ to homolog genes/proteins with similar sequences and marker trait associations (MTAs). 80% identity was used as a threshold in reporting putative homolog proteins.

## Acknowledgment

The research was conducted in the Feed and Forage Development (FFD) program at the ILRI forage genebank in Addis Ababa, Ethiopia, and the Earlham Institute, UK. The authors would like to thank The Royal Society, The Future Leaders – African Independent Research (FLAIR) Fellowships, FAPEMIG (Fundação de Amparo à Pesquisa do Estado de Minas Gerais - project APQ-03630-23), and the CGIAR intiative Sustainable Animal Productivity for Livelihoods, Nutrition and Gender inclusion initiative (SAPLING) for financial support. Embrapa, KALRO, Lanzhou University and USDA are acknowledged for making their germplasm and breeding lines available in this study. We acknowledge Collins Mutai for the DNA extraction of samples from Kenya. The authors would also like to thank Jane Poole for reading, commenting, and correcting the manuscript.

## Data availability statement

The data generated and presented in this study are deposited in the European Nucleotide Archive (ENA: https://www.ebi.ac.uk/ena/browser/view/PRJEB73794).

## Author contribution

TA, JDG and CJ designed and supervised the project and the manuscript writing, TA, JDG, LH and HJ analyzed phenotypic and genotypic datasets. AY, LH collected leaf samples and extracted DNA. TA, MSM, HE and NAT involved in the supervision of the phenotyping project in Ethiopia. TMO, ZJ, AB, NDO and QY involved in the supervision of the genotyping project. JFP, ALSA, and JCM and FJSL overssaw the field trial in Brazil. All authors have read, reviewed and approved this manuscript.

## Declarations

Ethics Approval and Consent to Participate

Not applicable.

## Consent for Publication

Not applicable.

---

**Supp. Fig 1.**
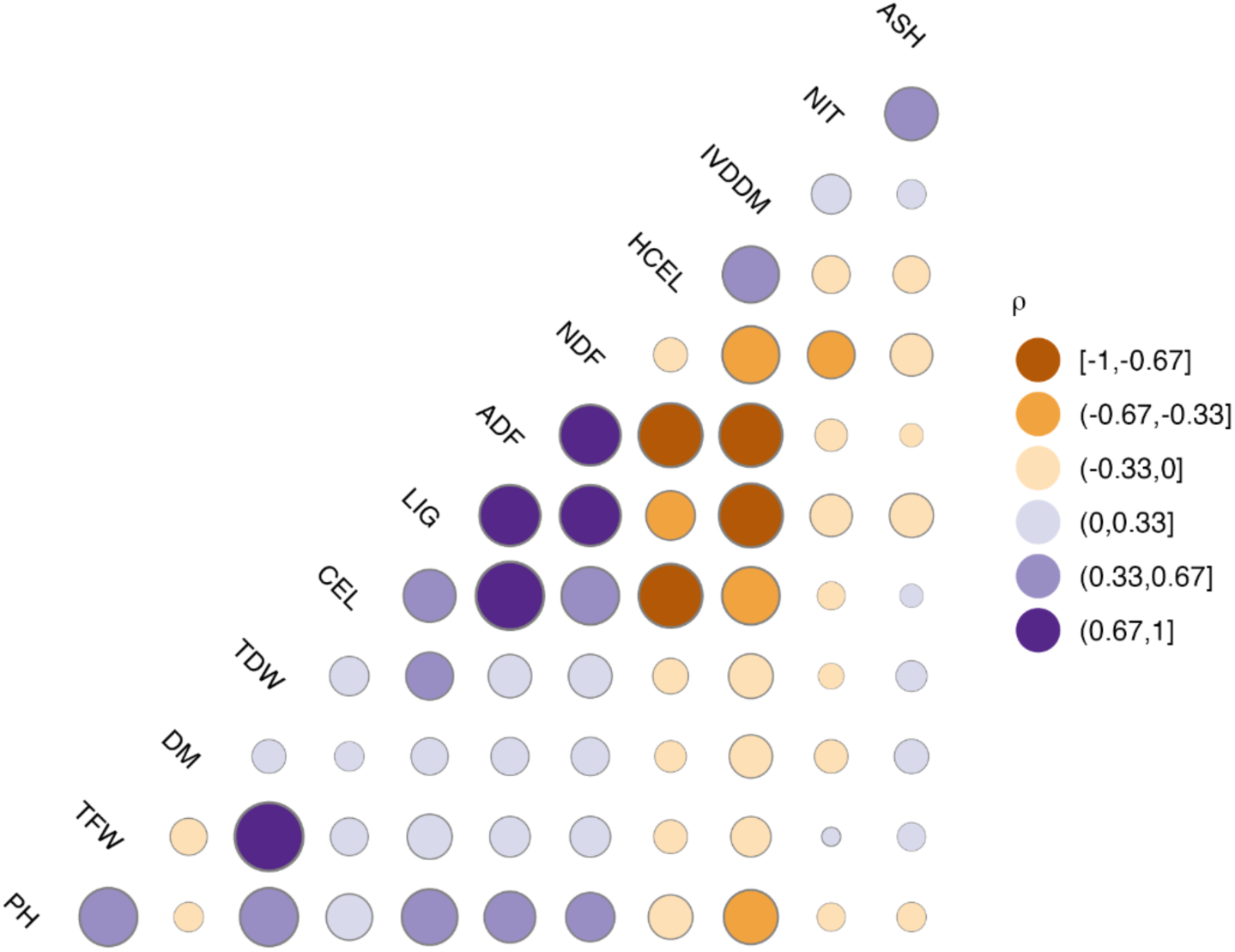
Correlation matrix plot of 12 quantitative traits measured in field trial in Brazil

**Supp. Fig 2.**
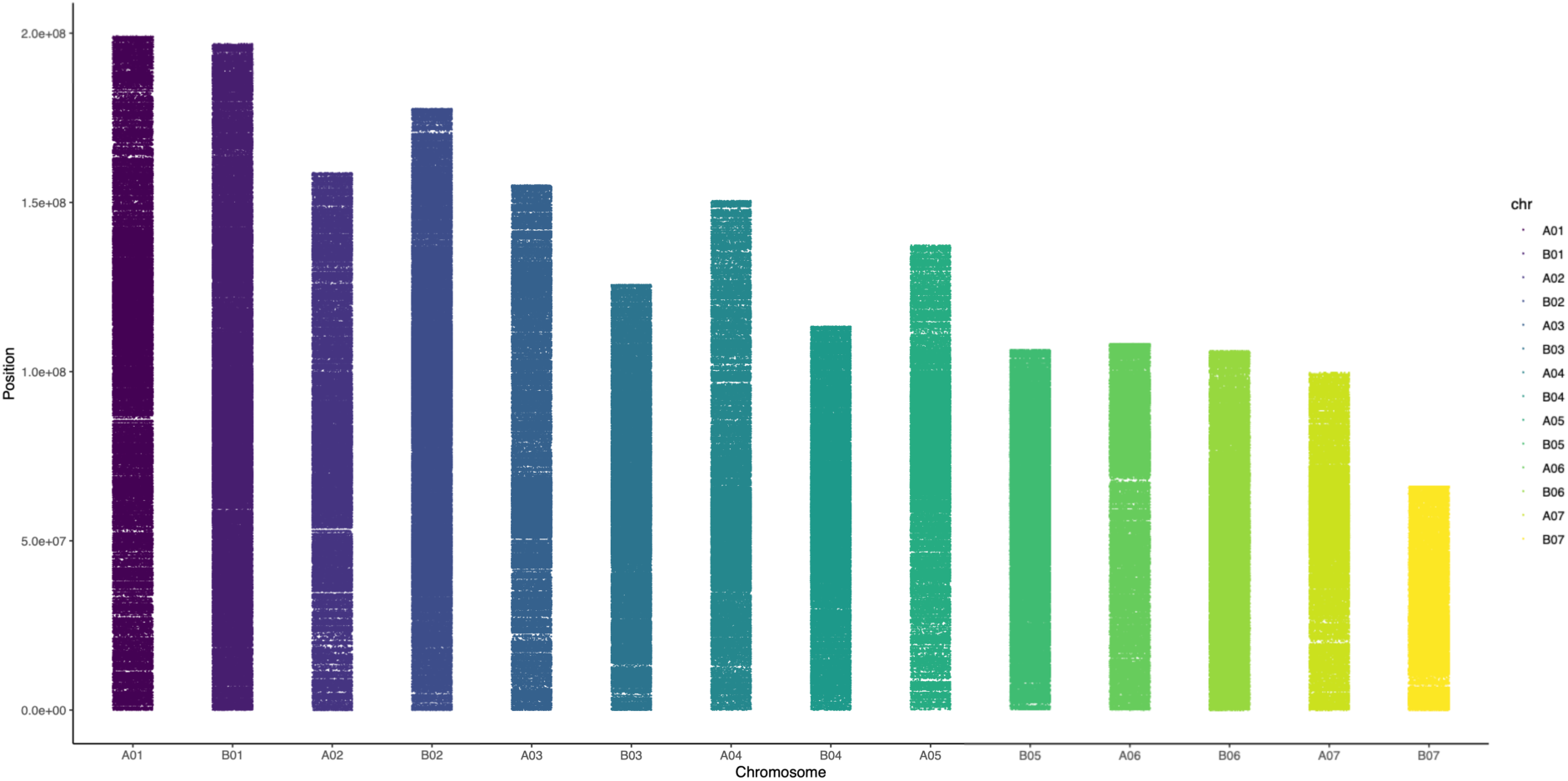
Distribution of half a million SNPs among 14 Napier grass chromosomes

**Supp. Fig. 3.**
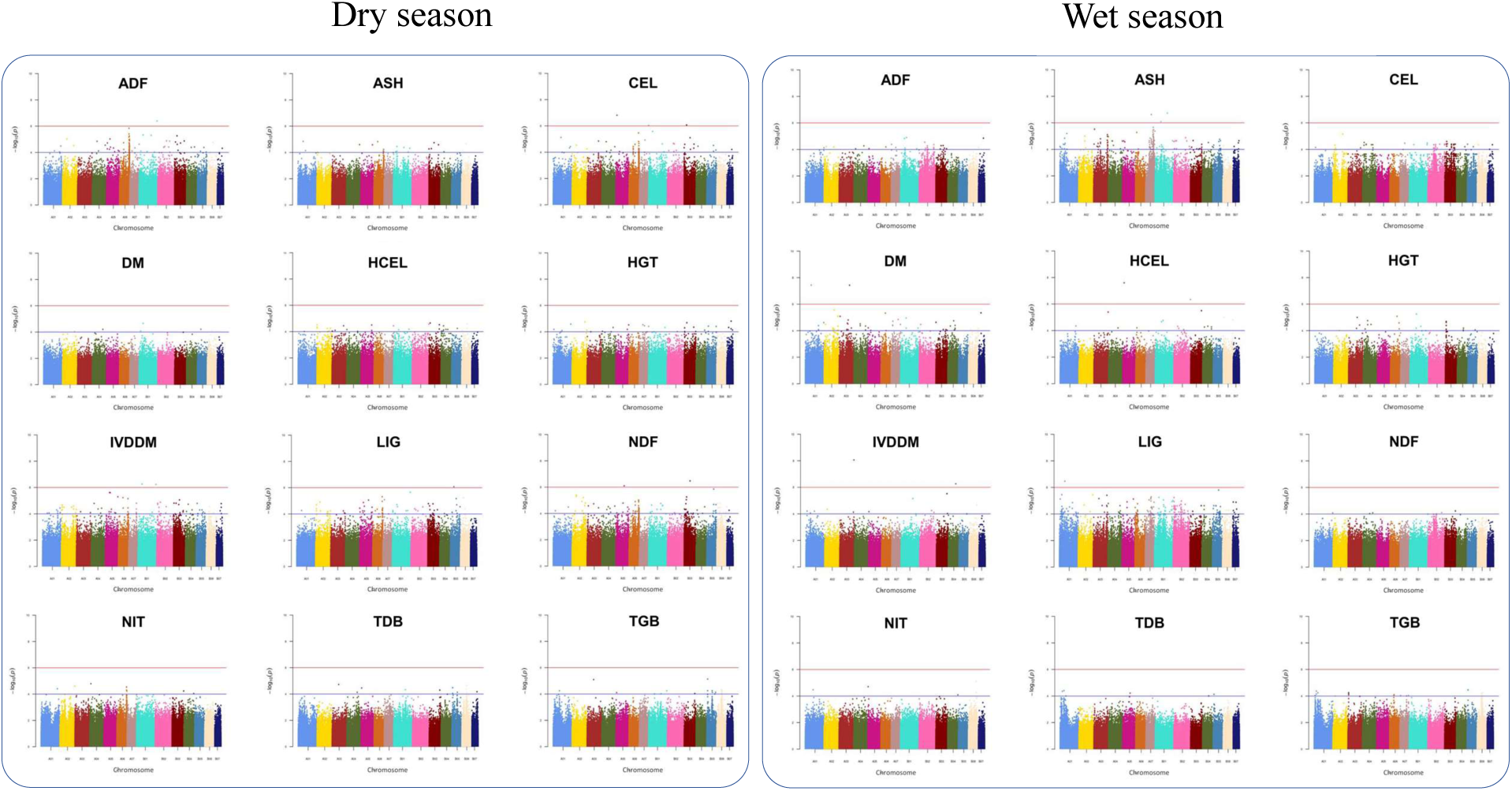
Significantly associated SNPs for the 12 quantitative traits evaluated during the dry (A) and wet (B) seasons in Brazil. Manhattan plots constructed with BLINK model. The GWAS analysis were performed for the following traits: Acid detergent fibre (ADF), ash (ASH), cellulose (CEL), dry matter concentration (DM), hemicellulose (HCEL), plant height (HGT), in vitro dry matter digestibility (IVDDM), lignin (LIG), neutral detergent fibre (NDF), nitrogen content (NIT), total dry weight (TDB), and total fresh weight (TGB). The blue and red lines represent the thresholds with p-value of 0.05 (–log10(p) > 4) and 0.01 (–log10(p) > 6), respectively

**Supp. Fig. 4.**
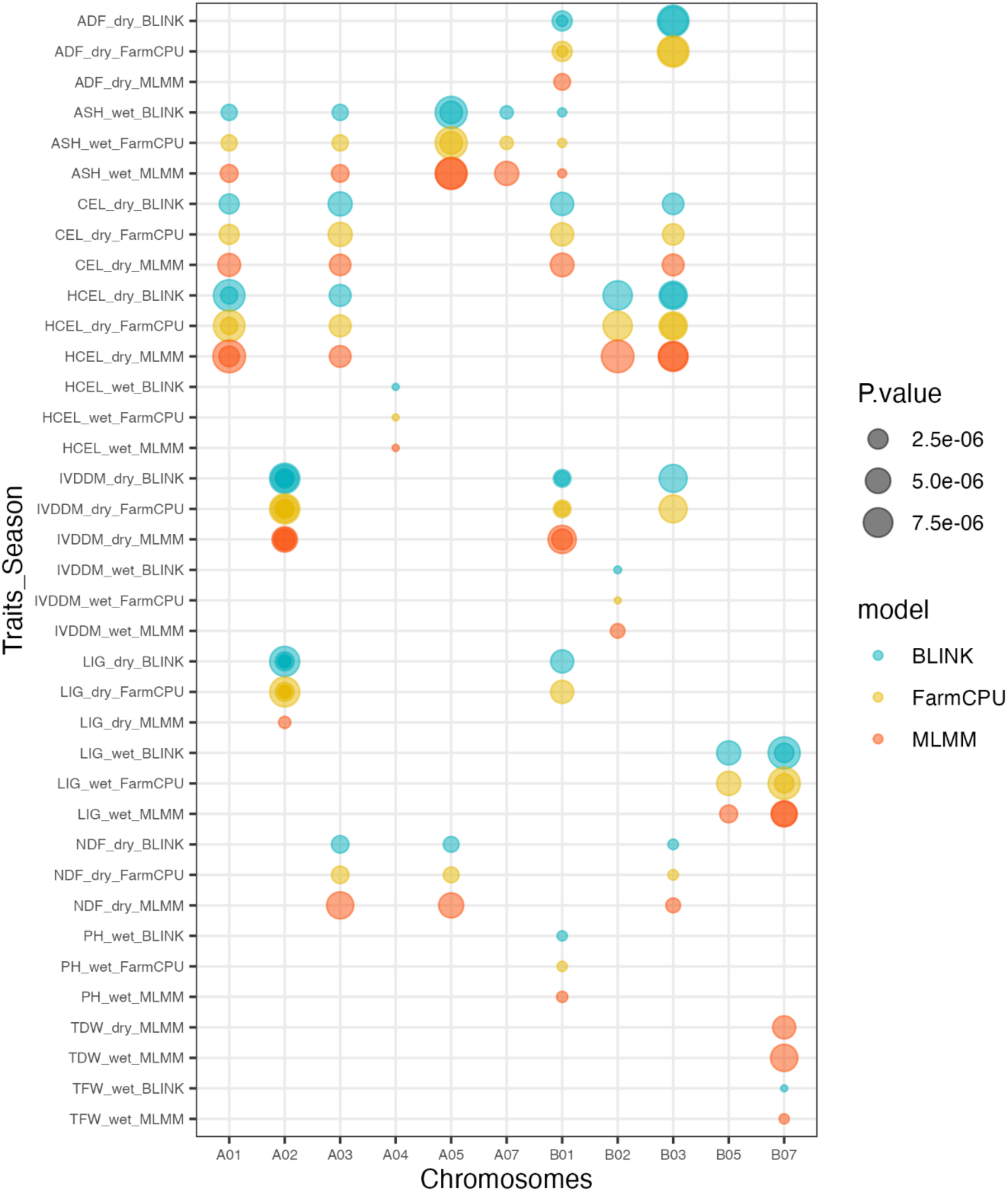
Bubble plot highlighting significantly associated SNPs (5 < −log10(p)) for 12 measured traits (dry and wet season) for the field trial in Brazil, with three GAPIT models

**Supp. Fig. 5.**
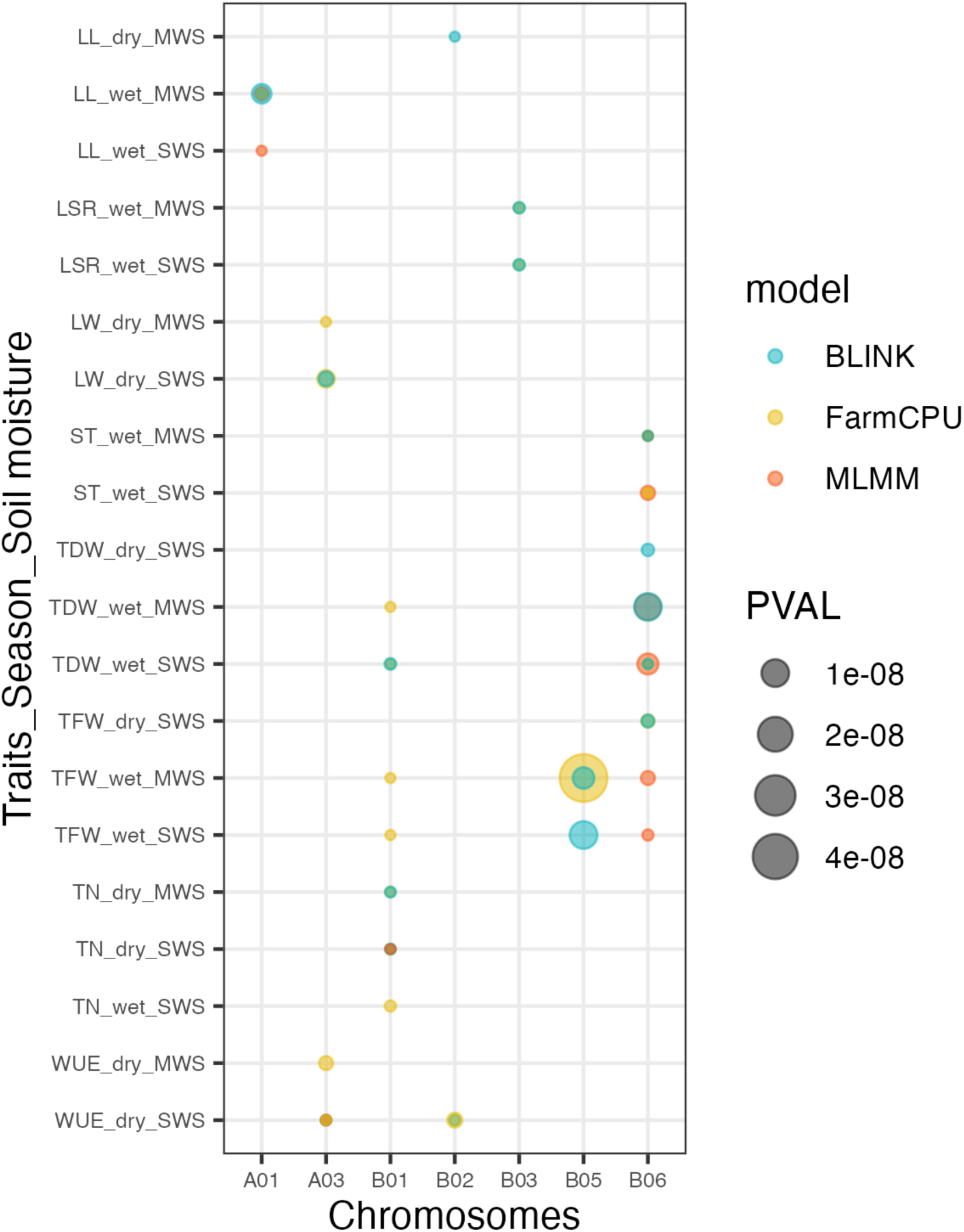
Bubble plot highlighting significantly associated SNPs (8 < −log10(p)) for 12 measured traits (dry and wet season) for the field trial in Ethiopia, with three GAPIT models

**Supp. Fig. 6.**
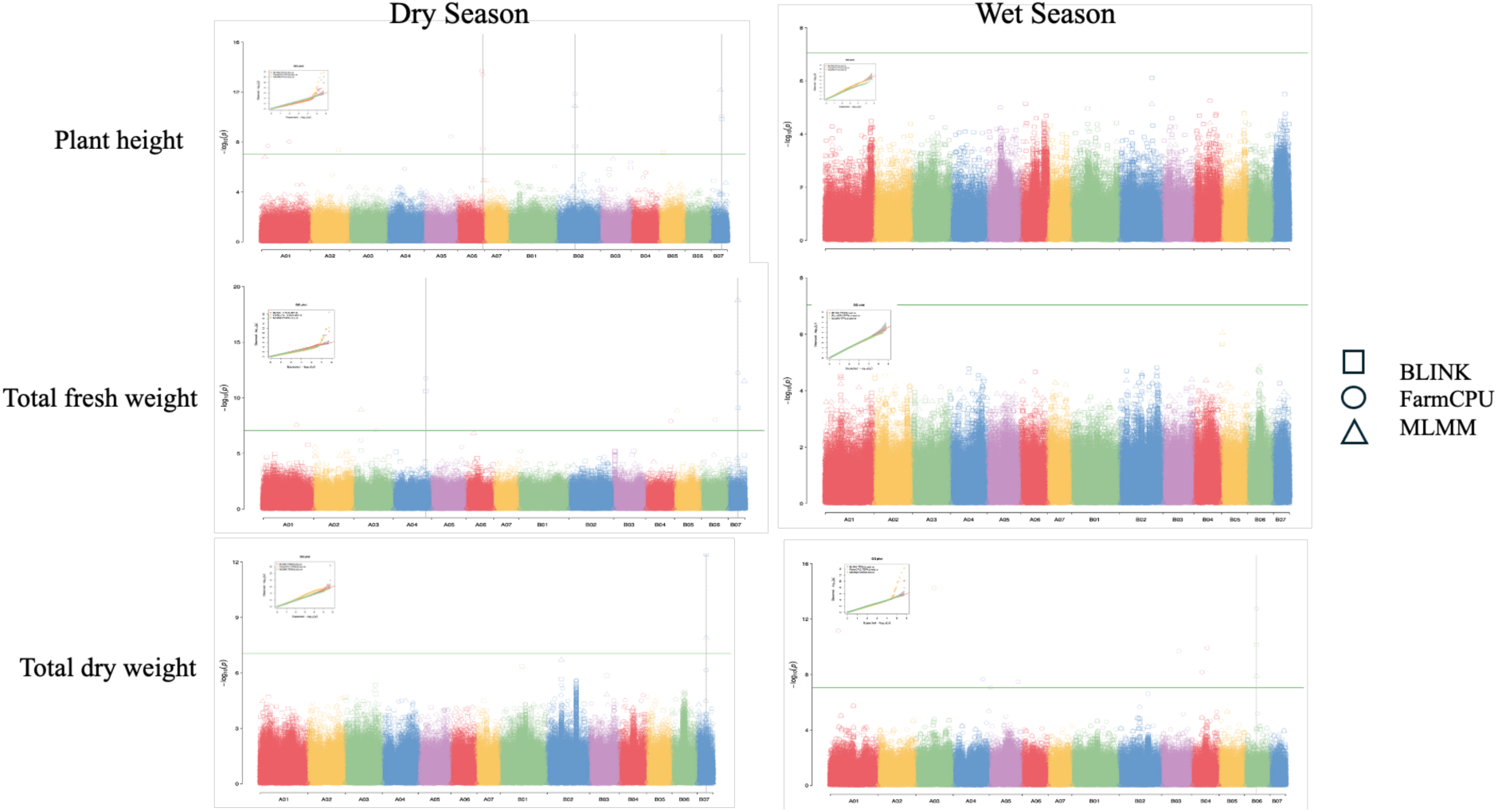
Manhattan plot of a genome-wide association study on three traits, plant height, total fresh weight and total dry weight with three GAPIT models. The genome-wide significance level is set at 7 × 10−8 and plotted as the dotted line. The most significant SNPs are above the aforementioned threshold. Quantile-quantile (QQ) plot for each analysis as also shown in the Manhattan plot. The different shapes correlated with three GAPIT models used

## References

1. FAO. Challenges and Opportunities for Carbon Sequestration in Grassland Systems: A Technical Report on Grassland Management and Climate Mitigation. (Rome: Food and Agriculture Organization of the United Nations), (2010).

2. Njuki, J. & Sanginga, P. C. “Gender and livestock: key issues, challenges and opportunities,” in Women, Livestock Ownership and Markets: Bridging the gender gap in Eastern and Southern Africa, eds J. Njuki and P. C. Sanginga (New York, NY: Routledge), (2013).

3. Simeão, R.M., Resende, M.D., Alves, R.S., Pessoa-Filho, M., Azevedo, A.L.S., Jones, C.S., Pereira, J.F., & Machado, J.C. Genomic selection in tropical forage grasses: current status and future applications. Frontiers in Plant Science, 12, 665195, (2021).

4. Balehegn, M., Kebreab, E., Tolera, A., Hunt, S., Erickson, P., Crane, T.A., & Adesogan, A.T., Livestock sustainability research in Africa with a focus on the environment. Animal Frontiers. (2021).

5. Balehegn, M., Duncan, A., Tolera, A., Ayantunde, A. A., Issa, S., Karimou, M., & Adesogan, A. T. Improving adoption of technologies and interventions for increasing supply of quality livestock feed in low-and middle-income countries. Global food security, 26, 100372 (2020).

6. Paul, B.K., Koge, J., Maass, B.L., Notenbaert, A., Peters, M., Groot, J.C., & Tittonell, P., Tropical forage technologies can deliver multiple benefits in Sub-Saharan Africa. A meta-analysis. Agronomy for Sustainable Development, 40 (4), 1–17 (2020).

7. Hanan, N.P., & Kahiu, M.N., Mapping Forage Resources Using Earth Observation Data: A Case Study to Assess the Relationship Between Herbaceous and Woody Cover Components as Determinants of Large Herbivore Distribution in Sub-Saharan Africa. In AGU Fall Meeting Abstracts, **Vol.** 2016, GC13D–1225 (2016).

8. Tolera, A. The role of forage supplements in smallholder mixed farming systems. In: Hare, M.D., Wongpichet, K. (Eds.), Forages: A Pathway to Prosperity for Smallholder Farmers, (Proceedings of an International Forage Symposium, Faculty of Agriculture. Ubon Ratchathani University, Thailand, 165–186 (2007).

9. Smith, J., Sones, K., Grace, D. MacMillan, S. Tarawali, S., & Herrero, M., Beyond milk, meat and eggs: Role of livestock in food and nutrition security. Animal Frontiers, 3 6–13 (2013).

10. Enahoro, D., Mason-D’Croz, D., Mul, M., Rich, K.M., Robinson, T.P. Thornton, P., & Staal, S.S. Supporting the sustainable expansion of livestock production in South Asia and Sub-Saharan Africa: Scenario analysis of investment options. Global food security; 20 114–121 (2019).

11. Mkutche, C.D. Evaluation of feed resources for local goat production under traditional management systems in Golomoti EPA Dedza and on-station at Bunda Campus, LUANAR, Malawi. Diss. International Institute of Tropical Agriculture, (2020).

12. Habte, E., Muktar, M.S., Abdena, A., Hanson, J., Sartie, A.M., Negawo, A.T., Machado, J.C., Ledo, F.J.D.S. & Jones, C.S. Forage performance and detection of marker-trait associations with potential for Napier grass (*Cenchrus purpureus*) improvement. Agronomy, 10 (4), 542 (2020).

13. Mwendia, S.W., Wanyoike, M., Nguguna, J.G.M., Wahome, R.G., & Mwangi, D.M. Evaluation of napier grass cultivars for resitance for Napier head smut. *Kenya Agricultural Research Institute*, University of Nairobi, (2000).

14. G. Simons, S. A. & Hillocks, R.J. Pests, diseases, and weeds of Napier grass, Pennisetum purpureum: a review. I. J. Pest Management, 48 (1), 39–48 (2002).

15. Orodho, A.B. The role and importance of Napier grass in the smallholder dairy industry in Kenya (2006).

16. Mengistu, A., Kebede, G., Feyissa, F. & Assefa, G. Review on major feed resources in Ethiopia: Conditions, challenges and opportunities. Academic Research Journal of Agricultural Science and Research, 5 (3), 176–185 (2017).

17. Maleko, D., Mwilawa, A., Msalya, G., Pasape, L., & Mtei, K. Forage growth, yield, and nutritional characteristics of four varieties of Napier grass (*Pennisetum purpureum Schumach*) in the west Usambara Highlands, Tanzania. Scientific African, 6, e00214 (2019).

18. Muyekho, F. 2015. Napier grass feed resource: Production, constraints, and implications for smallholder farmers in East and Central Africa, (2015).

19. Muktar, M.S., Habte, E., Teshome, A., Assefa, Y., Negawo, A.T., Lee, K.W., Zhang, J., & Jones, C.S. Insights into the Genetic Architecture of Complex Traits in Napier Grass (*Cenchrus purpureus*) and QTL Regions Governing Forage Biomass Yield, Water Use Efficiency and Feed Quality Traits. Frontiers in plant science, 12, 678862 (2022).

20. Anderson, W. F., Dien, B. S., Brandon, S. K., & Peterson, J. D. Assessment of bermudagrass and bunch grasses as feedstock for conversion to ethanol. Applied biochemistry and biotechnology, 145 (1), 13–21 (2008).

21. Knoll, J.E. & Anderson, W.F. Vegetative propagation of Napiergrass and energy cane for biomass production in the Southeastern United States. Agronomy Journal, 104 2, 518–522 (2012).

22. Rocha, J.R., D.A.S.C., Marçal, T.S., Salvador, F.V., da Silva, A.C., Carneiro, P.C.S., de Resende, M.D.V., Carneiro, J.D.C., Azevedo, A.L.S., Pereira, J.F., & Machado, J.C. Unraveling candidate genes underlying biomass digestibility in elephant grass (*Cenchrus purpureus*). BMC Plant Biol. 19 1, 548, (2019).

23. Dokbua, B., Waramit, N., Chaugool, J., & Thongjoo, C. Biomass Productivity, Developmental Morphology, and Nutrient Removal Rate of Hybrid Napier Grass (*Pennisetum purpureum x Pennisetum americanum*) in Response to Potassium and Nitrogen Fertilization in a Multiple-Harvest System. Bioenergy Research, 14 (4), 1106–1117 (2021).

24. Fukagawa, S., & Isshi, Y. Grassland Establishment of Dwarf Napier grass (*Pennisetum purpureum* Schumach) by Planting of Cuttings in the Winter Season. Agronomy. 8 12, 1–10 (2018).

25. Yan, Q., Wu, F. Xu, P., Sun, Z., Li, J., Gao, L., Lu, L., Chen, D., Muktar, M., Jones, C., & Yi, X. The elephant grass (*Cenchrus purpureus*) genome provides insights into anthocyanidin accumulation and fast growth. Molecular ecology resources, 21 (2), 526–542 (2021).

26. Zhang, S., Xia, Z., Li, C., Wang, X., Lu, X., Zhang, W., Ma, H., Zhou, X., Zhang, W., Zhu, T., Liu, P., Liu, G., Wang, W., & Xia, T. Chromosome-scale genome assembly provides insights into speciation of allotetraploid and massive biomass accumulation of elephant grass (*Pennisetum purpureum Schum*.). Mol Ecol Resour. 22 6:2363–2378, (2022).

27. Kamau, M. Farm household allocative efficiency: a multi-dimensional perspective on labour use in Western Kenya. Wageningen University and Research; (2007).

28. Habte, E., Teshome, A., Muktar, M.S., Assefa, Y., Negawo, A.T., Machado, J.C., Ledo, F.J.D.S., & Jones, C.S. Productivity and Feed Quality Performance of Napier Grass (*Cenchrus purpureus*) Genotypes Growing under Different Soil Moisture Levels. Plants, 11 19, 2549 (2022).

29. Lamb, M.C., Anderson, W.F., Strickland, T.C., Coffin, A.W., Sorensen, R.B., Knoll, J.E., & Pisani, O. Economic competitiveness of Napier grass in irrigated and non-irrigated Georgia coastal plain cropping systems. BioEnergy Research, 11 574–582, (2018).

30. O’Brien, D. & J. Suszkiw. Finding the right biofuels for the south-east: A range of alternatives. Agric. Res. 60 10, (2012).

31. Sawasdee, V. & Pisutpaisal, N. Potential of Napier grass Pak Chong 1 as feedstock for biofuel production. Energy Reports, 7, 519–526, (2021).

32. Paudel, D., Kannan, B., Yang, X., Harris-Shultz, K., Thudi, M., Varshney, R.K., Altpeter, F. & Wang, J. Surveying the genome and constructing a high-density genetic map of napiergrass (*Cenchrus purpureus* Schumach). Scientific reports, 8 (1), 1–11 (2018).

33. Muktar, M. S., Teshome, A., Hanson, J., Negawo, A. T., Habte, E., & Domelevo Entfellner, J. B., et al. Genotyping by sequencing provides new insights into the diversity of Napier grass (*Cenchrus purpureus*) and reveals variation in genome-wide LD patterns between collections. Sci. Rep. 9, 1–15 (2019).

34. Muktar, M.S., Bizuneh, T., & Anderson, W., et al. Analysis of global Napier grass (*Cenchrus purpureus*) collections reveals high genetic diversity among genotypes with some redundancy between collections. Scientific reports, 13, 14509 (2023).

35. Gupta, S. C., & Mhere, O. Identification of superior pearl millet by napier hybrids and Napier’s in Zimbabwe. African Crop Sci. J. 5, 229–237 (1997).

36. Wanjala, B.W., Obonyo, M., Wachira, F.N., Muchugi, A., Mulaa, M., Harvey, J., Skilton, R.A., Proud, J., & Hanson, J. Genetic diversity in Napier grass (*Pennisetum purpureum*) cultivars: implications for breeding and conservation. AoB PLANTS 5 plt022 (2013).

37. Pereira, A.V., Lédo, F. J., da, S. & Machado, J. C. BRS Kurumi and BRS Capiaçu - New elephant grass cultivars for grazing and cut-and-carry system. Crop Breeding and Applied Biotechnology, 17, 59–62, (2017).

38. Negawo, A.T., Jorge, A., Hanson, J., Teshome, A., Muktar, M.S., Azevedo, A.L.S., Lédo, F.J.d.S., Machado, J.C. & Jones, C.S. Molecular markers as a tool for germplasm acquisition to enhance the genetic diversity of a Napier grass (*Pennisetum purpureum*) collection. Tropical Grasslands 6 (2):58–69 (2018).

39. Martel, E., De Nay, D., Siljak-Yakovlev, S., Brown, S., & Sarr, A. Genome size variation and basic chromosome number in pearl millet and fourteen related Pennisetum species. J. Hered. 88 139–143 (1997).

40. Hanna, W.W., Chaparro, C.J., Mathews, B.W., Burns, J.C., Sollenberger, L.E., & Carpenter, J.R. Perennial pennisetums. Warm-season (C4) grasses; 45:503–35 (2004).

41. Foster, C.A. Interpopulation and intervarietal hybridization in Lolium perenne breeding: heterosis under non-competitive conditions. J. Agric. Sci. 76, 107–130 (1971).

42. Pembleton, L. W., Wang, J., Cogan, N. O. I., Pryce, J. E., Ye, G., & Bandaranayake, C. K., et al. Candidate gene-based association genetics analysis of herbage quality traits in perennial ryegrass (*Lolium perenne* L.). Crop Pasture Sci. 64, 244–253, (2013).

43. Rengsirikul, K., Ishii, Y., Kangvansaichol, K., Sripichitt, P., Punsuvon, V., Vaithanomsat, P., Nakamanee, G., & Tudsri, S. Biomass yield, chemical composition, and potential ethanol yields of 8 cultivars of napiergrass (Pennisetum purpureum Schumach.) harvested 3-monthly in central Thailand, (2013).

44. Tessema, B.B., & Liu, H., Sørensen, A.C. Andersen, J.R. Jensen, J. Strategies Using Genomic Selection to Increase Genetic Gain in Breeding Programs for Wheat. Front. Genet. 11 578123 (2020).

45. Camargo, L.S.A., & Pereira, J.F. Genome-editing opportunities to enhance cattle productivity in the tropics. CABI Agriculture and Bioscience, 3 1, 8 (2022).

46. Kruger, M., Davies, N., Myburgh, K., & Lecour, S. Proanthocyanidins, anthocyanins and cardiovascular diseases. Food Research International, 59, 46, (2014).

47. Yao, N., Xian-Feng, Y. I., Lai, Z. Q., Liang, Y. L., Deng, S. Y., & Lai, D.W. Effects of Pennisetum purpureum Schumab cv. Purple on growth performance and serum biochemical parameters of meat geese. Journal of Southern Agriculture, 47 (12), 2163–2168, (2016).

48. Hardie, D.G. Plant protein serine/threonine kinases: classification and functions. Annual review of plant biology, 50(1), 97–131 (1999).

49. Jose, J., Ghantasala, S. & Roy, C. S., Arabidopsis transmembrane receptor-like kinases (RLKs): a bridge between extracellular signal and intracellular regulatory machinery. International Journal of Molecular Sciences, 21(11), 4000 (2020).

50. Liu, J., Li, W., Wu, G. & Ali, K., An update on evolutionary, structural, and functional studies of receptor-like kinases in plants. Frontiers in Plant Science, 15, p.1305599. (2024).

51. Shumskaya M & Wurtzel ET. The carotenoid biosynthetic pathway: thinking in all dimensions. Plant Science. (2013).

52. Stanley, L. & Yuan, Y. Transcriptional Regulations of carotenoid biosynthesis in plants:so many regulators, so little consensus. Frontiers in Plant Science, 10, (2019).

53. Abdelali H. & Zakir H. Carotenoid Biosynthesis and Regulation in Plants, Agriculture and Agri-Food Canada, 1391 Sandford Street, London, Ontario, Canada, (2016).

54. Peterson, R.A. bestNormalize: Normalizing transformation functions. R package version. 1 (573 (2018).

55. R Core Team R: A language and environment for statistical computing. R Foundation for Statistical Computing, Vienna, Austria., URL https://www.R-project.org/ (2022)

56. Kassambara, A. Practical guide to principal component methods in R: PCA, M (CA), FAMD, MFA, HCPC, factoextra. Sthda; (2017).

57. Andrews, S. FastQC: A Quality Control Tool for High Throughput Sequence Data. http://www.bioinformatics.babraham.ac.uk/projects, (2010).

58. Ewels, P., Magnusson, M., Lundin, S., & Käller, M. MultiQC: summarize analysis results for multiple tools and samples in a single report. Bioinformatics; 32 (19), 3047–3048 (2016).

59. Bolger, A.M., Lohse, M., & Usadel, B. Trimmomatic: a flexible trimmer for Illumina sequence data. Bioinformatics, 30 (15), 2114–2120 (2014).

60. Li, H. & Durbin, R. Fast and accurate short read alignment with Burrows-Wheeler transform. Bioinformatics, 25 (14),1754–1760 (2009).

61. Li, H., Handsaker, B., Wysoker, A., Fennell, T., Ruan, J., & Homer, N. et al. The sequence alignment/map format and SAMtools. Bioinformatics 25, 2078–2079 (2009).

62. McKenna, A., Hanna, M. & Banks, E. et al. The Genome Analysis Toolkit: A MapReduce framework for next-generation DNA sequencing data. Genome research 20;1297–303 (2010).

63. Alexander, D. H., & Lange, K. Enhancements to the ADMIXTURE algorithm for individual ancestry estimation. BMC Bioinform, 12 (1):246 (2009).

64. Jakobsson, M., & Rosenberg, N.A (CLUMPP: a cluster matching and permutation program for dealing with label switching and multimodality in analysis of population structure. Bioinformatics 23 (14):1801–1806 (2007).

65. Francis R.M. pophelper: an R package and web app to analyze and visualize population structure. Mol ecol res. 17 (1), 27–32 (2017).

66. Zheng, X., Levine, D., Shen, J., Gogarten, S.M., Laurie, C., & Weir, B.S. A high-performance computing toolset for relatedness and principal component analysis of SNP data. Bioinformatics, 28 (24), 3326–8, (2012).

67. Sievert, C. Interactive Web-Based Data Visualization with R, plotly, and shiny. CRC Press. ISBN: 9781138331457, https://plotly-r.com. (2020).

68. Letunic I., & Bork P. Interactive tree of life (iTOL) v5: An online tool for phylogenetic tree display and annotation. Nucleic Acids Res. 49 (W1): W293–6. (2021).

69. Lipka, A.E., Tian, F., Wang, Q., & Peiffer, J. Li, M., Bradbury, P. J., Gore, M.A., Buckler, E.S., & Zhang Z. GAPIT: genome association and prediction integrated tool. Bioinformatics.; 28 (18); 2397–9, (2012).

70. Huang, M., Liu, X., Zhou, Y., Summers, R.M., & Zhiwu Z. BLINK: a package for the next level of genome-wide association studies with both individuals and markers in the millions. GigaScience 28 (18), (2019).

71. Wang, J., & Zhang, Z. GAPIT Version 3: Boosting Power and Accuracy for Genomic Association and Prediction. Genom. Proteom Bioin. 629–640 (2021).

72. Goodstein, D.M., Shu, S., Howson, R., Neupane, R., Hayes, R. D., Fazo, J., Mitros, T., Dirks, W., Hellsten, U., Putnam, N. & Rokhsar, DS. Phytozome: a comparative platform for green plant genomics. Nucleic acids research.;40 (D1): D1178–86 (2012).

